# Bone regeneration in Ds-Red pig calvarial defect using allogenic transplantation of EGFP-pMSCs – a comparison of host cells and seeding cells in the scaffold

**DOI:** 10.1101/599670

**Authors:** Ming-Kai Hsieh, Chia-Jung Wu, Xuan-Chun Su, Yi-Chen Chen, Tsung-Ting Tsai, Chi-Chien Niu, Po-Liang Lai, Shinn-Chih Wu

**Affiliations:** Institute of Biotechnology, National Taiwan University, Taipei, Taiwan; Department of Orthopaedic Surgery, Chang Gung Memorial Hospital, Taoyuan, Taiwan; Bone and Joint Research Center, Chang Gung Memorial Hospital, Taoyuan, Taiwan; College of Medicine, Chang Gung University, Taoyuan, Taiwan; Department of Animal Science and Technology, National Taiwan University, Taipei, Taiwan; Center for Biotechnology, National Taiwan University, Taipei, Taiwan

**Keywords:** mesenchymal stem cell, bone regeneration, enhanced green fluorescent, DsRed pig, pig calvarial defect model

## Abstract

**Background:** Cells, scaffolds, and factors are the triad of regenerative engineering; however, it is difficult to distinguish whether cells in the regenerative construct are from the seeded cells or host cells via the host blood supply. We performed a novel *in vivo* study to transplant enhanced green fluorescent pig mesenchymal stem cells (EGFP-pMSCs) into calvarial defect of DsRed pigs. The cell distribution and proportion were distinguished by the different fluorescent colors through the whole regenerative period.

**Method/Results:** Eight adult domestic Ds-Red pigs were treated with five modalities: empty defects without scaffold (group 1); defects filled only with scaffold (group 2); defects filled with osteoinduction medium-loaded scaffold (group 3); defects filled with 5 × 10^3^ cells/scaffold (group 4); and defects filled with 5 × 10^4^ cells/scaffold (group 5). The *in vitro* cell distribution, morphology, osteogenic differentiation, and fluorescence images of groups 4 and 5 were analyzed. Two animals were sacrificed at 1, 2, 3, and 4 weeks after transplantation. The *in vivo* fluorescence imaging and quantification data showed that EGFP-pMSCs were represented in the scaffolds in groups 4 and 5 throughout the whole regenerative period. A higher seeded cell density resulted in more sustained seeded cells in bone regeneration compared to a lower seeded cell density. Host cells were recruited by seeded cells if enough space was available in the scaffold. Host cells in groups 1 to 3 did not change from the 1st week to 4th week, which indicates that the scaffold without seeded cells cannot recruit host cells even when enough space is available for cell ingrowth. The histological and immunohistochemical data showed that more cells were involved in osteogenesis in scaffolds with seeded cells.

**Conclusion:** Our *in vivo* results showed that more seeded cells recruit more host cells and that both cell types participate in osteogenesis. These results suggest that scaffolds without seeded cells may not be effective in bone transplantation.

## Introduction

Skeletal defects require surgery using bone grafts. Autografts are the gold standard for bone grafting [1]; however, donor site morbidity and the limited amount of available donor tissue restrict their application [2, 3]. Regenerative tissue engineering using cells, scaffolds, factors and blood supply [4] has become an alternative method to treat skeletal bone defects.

Allografts may provide the same osteoconductive conduit for bony fusion as traditional autografts and may have comparable biomechanical properties without amount restriction [5, 6]. Although depleted of osteoprogenitor cells like mesenchymal stem cells (MSCs), the fusion rate still reaches 73% to 100% in instrumented spinal fusion [7–16], making allograft a clinically feasible alternative form of fusion. The role of cells in allograft prompted us to wonder if seeded cells are necessary in bone regeneration clinically.

MSCs are undifferentiated multipotent cells with the capacity to differentiate into osteoblasts, chondrocytes, adipocytes, fibroblasts, and other tissues of mesenchymal origin [17]. MSCs lack expression of costimulatory molecules like CD40, CD80, and CD86, which makes them largely non-immunogenic [18], as supported by evaluations of their immunosuppressive properties using mitogen proliferation assays [19]. Due to these suitable transplantation properties, MSCs are an important material in developmental biology and transgenic methods, as well as in potential clinical applications in tissue engineering and gene therapy [20, 21].

Some studies have found that seeded MSCs may be capable of releasing major growth factors, reducing the immune response, mobilizing the host’s cells or eventually directly differentiating into osteoblasts [22-25]. The expansion, proliferation, migration, viability and osteogenic differentiation of MSCs on different types of scaffolds have been shown in studies [26-31], but the role of seeded MSCs is still poorly addressed. Histologically, it is difficult for us to distinguish cells in the regenerative construct from seeded MSCs or from host MSCs via the host blood supply. In this study, we transplanted EGFP pig MSCs into calvarial defect of DsRed pigs. The cell distribution and proportion were distinguished by different fluorescent colors throughout the whole regenerative period.

The purpose of the present study was to clarify the distribution and proportion of seeded cells and host cells by tracking two fluorescent cells in the same scaffold in a pig critical-sized calvarial defect model.

## Materials and methods

### EGFP-pMSCs culture in the scaffold

#### Scaffold

A hemostatic gelatin sponge, Spongostan^TM^ (Ferrosan Medical Device, MS0003, thickness 0.1 cm), was used as the 3D scaffold [32]. Scanning electron microscopy (SEM) (Hitachi, SU8220) analysis indicated a mean pore size of approximately 148 ± 62 μm (Fig 1). To perform transplantation, the scaffolds were cut into disks with a diameter of 0.8 cm, sterilized by 75 % (v/v) ethanol and washed three times with phosphate-buffered saline. The sterile scaffold disks were then immersed in Opti-MEM medium (Gibco) before use.

**Fig 1.**
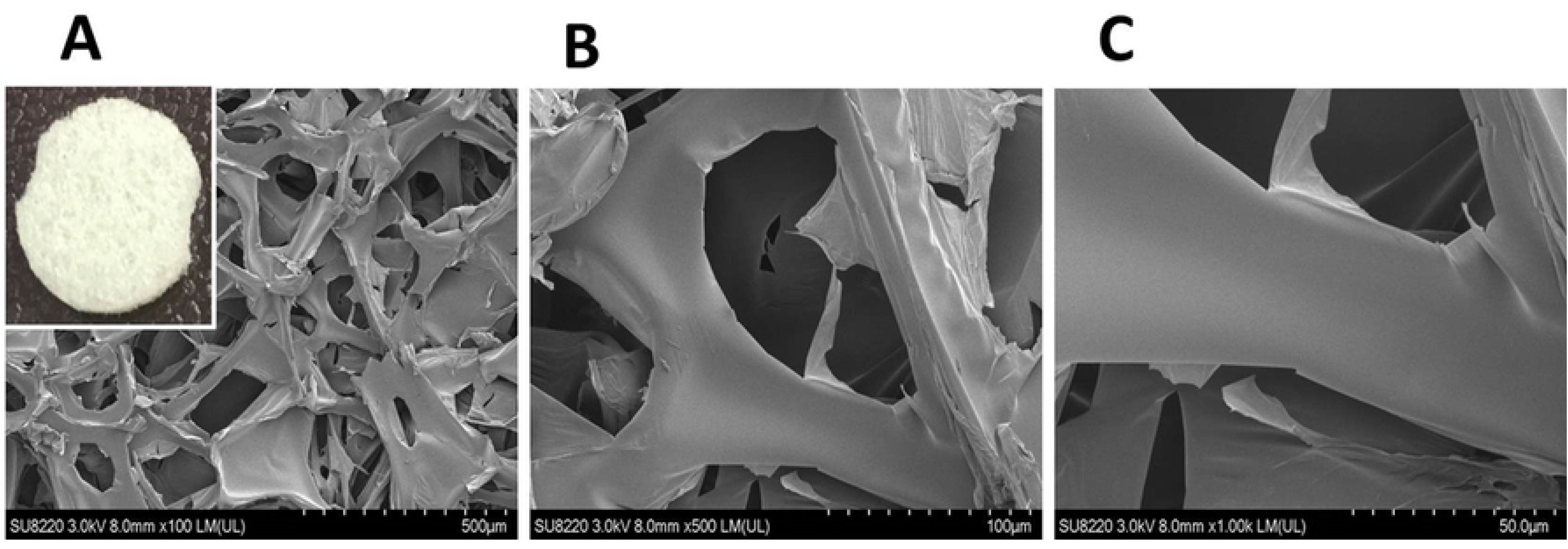
Morphology of the scaffold. The Hemostatic Gelatin Sponge, SpongostanTM (Ferrosan Medical Device, MS0003, Thickness 0.1 cm) was used as the 3D Scaffold and determined by SEM (Scanning Electron Microscopy) to have a mean pore size of approximately 148 ± 62 μm. Increased magnification was observed from A to C.

#### Cell culture

Passage 5 EGFP-pMSCs [19] were cultured in minimum essential medium alpha (MEM-α) (Gibco) supplemented with 10% fetal bovine serum (HyClone) and antibiotic solutions (100 U/mL penicillin, 100 *μ*g/mL streptomycin, 50 *μ*g/mL gentamicin and 250 ng/mL fungizone) (Gibco). The cells were maintained in a humidified 37 ℃ incubator supplied with 5 % CO_2_.

#### Seeding of cells

The sterile scaffold disks were individually placed onto the wells of a 48 well-plate. The cell seeding density was 5 × 10^3^ or 5 × 10^4^ cells/disk. Cells in 50 *μ*L of medium were seeded throughout the surface of the scaffold disk and incubated for 3 hours to allow cell attachment before the addition of medium. The culture condition was observed by fluorescent microscopy and recorded until the 28^th^ day after seeding onto scaffolds.

### *In vitro* cell distribution and morphology analysis

The cell scaffold constructs were washed twice with phosphate buffered saline (PBS; GIBCO™, Invitrogen Corp., Carlsbad, CA) and fixed in 1.5% v/v glutaraldehyde in 0.14M sodium cacodylate (pH 7.4) for 30 mins at room temperature. Dehydration was performed by sequential immersion in serial diluted ethanol solutions of 50, 60, 70, 80, 90, and 100% v/v. The samples were then transferred to hexamethyldisilazane and air-dried at room temperature overnight. The cell distribution and morphology were analyzed by SEM and confocal laser scanning microscopy (CLSM) (Bio-Rad MRC 600).

### *In vitro* osteogenic differentiation

The above two groups then received 0.5 mL/well of osteogenic induction medium (OIM) to promote cell differentiation. The OIM medium was composed of complete MEM-α medium enriched with 10^−7^ M dexamethasone (Sigma), 10 mM *β*-glycerophosphate (Sigma) and 50 *μ*g/mL ascorbic acid (Sigma) [32]. The OIM medium was replenished every two or three days for a total of 7 days. The osteogenic potential was assessed using an alkaline phosphatase staining and Alizarin red S (ARS) staining.

### Alkaline phosphatase staining and quantification

The osteogenic potential was assessed using an ALP assay kit (BioVision K412-500) after 7 days. The cells were washed twice with ice-cold PBS and lysed using 300 μL of RIPA (radioimmunoprecipitation assay) lysis buffer (Sigma-Aldrich Corp., St Louis, MO, USA) for 5 mins on ice. The cells were then rapidly scraped from the plate, and the cell lysates/RIPA buffer were transferred to a 1.5-mL microcentrifuge tube on ice for 20 min, followed by centrifugation at 8000 g for 10 min at 4 °C. The supernatant was then added to a new 1.5-mL microcentrifuge tube and stored at −20 °C. 50 µL of 5 mM pNPP solution was added to each well containing the test samples and the aliquots were incubated for 60 min at 25 °C while protected from light. Subsequently, 20 µL of the stop solution was added to terminate the ALP activity in the sample. The absorbance at 405 nm was measured with a spectrophotometer (UV-Vis 8500). Then, the ALP activity was calculated using the following formula: ((optical density –mean optical density of the control wells) × total volume × dilution)/(18.45 × sample volume).

### Alizarin red S (ARS) staining and quantification

Cells cultured on the discs at different time points were washed twice with PBS and fixed with 4% paraformaldehyde for 15 min. The fixative was removed, and the discs with cells were rinsed with deionized water. The scaffolds were stained with a 2% ARS staining kit (ScienCell) for 5 mins to clarify the calcium deposits. The staining solution was discarded, and the scaffolds were carefully washed with deionized water to remove excess stain. The staining results were captured with a digital camera. To quantify the staining of each scaffold, 1 mL/disc of 10 wt% cetylpyridinium chloride (Sigma) was added to the discs. The discs were left on an orbital shaker (speed, 60 rpm) for 1 h to completely resolve the dye from the discs. Finally, 100 μL of the dissolved solution from each scaffold was placed into a well of a 96-well plate, and the absorbance was measured at 540 nm with an ELISA reader.

### *In vivo* experimental design

Eight adult domestic Ds-Red pigs (average age of 15 ± 3months) with a mean weight of 100 ±20 kgs were obtained from the Department of Animal Science, National Taiwan University (Taipei, Taiwan). This study was carried out in strict accordance with the recommendations in the *Guide for the Care and Use of Laboratory Animals* of the National Institutes of Health. All animals were used under approved animal protocols by the Institutional Animal Care and Use Committee of National Taiwan University (NTU107-EL-00128) and housed accordingly.

The five treatment modalities were empty defects without scaffold (group 1; n=1); defects filled with only scaffold (group 2; n=1); defects filled with osteoinduction medium-loaded scaffold (group 3; n=1); defects filled with 5 × 10^3^ cells/scaffold (group 4; n=2) and defects filled with 5 × 10^4^ cells/scaffold (group 5; n=2).

Seven defects were created in each animal. The treatment modalities of the defects were determined for each animal separately by a randomization chart.

### Operation

Zoletil^®^ (5 mg/kg body weight) was injected in the neck area of the pig intramuscularly as a sedative. All experimental procedures were performed under a standard anesthetic/ analgesic protocol as described in previous studies [19, 33].

The frontal bone was exposed after a standard sagittal approach in the forehead region. Identical bony defects were then created with a saline-cooled trephine burr (diameter 8 mm, depth 2 mm) meeting the requirements for a critical size defect in pigs [34]. The defects were positioned at least 5 mm apart to minimize biological interactions [35]. The internal plate of the neurocranium remained completely intact during the procedure. After well hemostasis, the scaffold was implanted (Fig 2). The periosteum and skin over the defects were sutured in two layers with resorbable material (Vicryl USP 3-0; Vicryl USP1-0; Ethicon Johnson & Johnson, NJ, USA).

**Fig 2.**
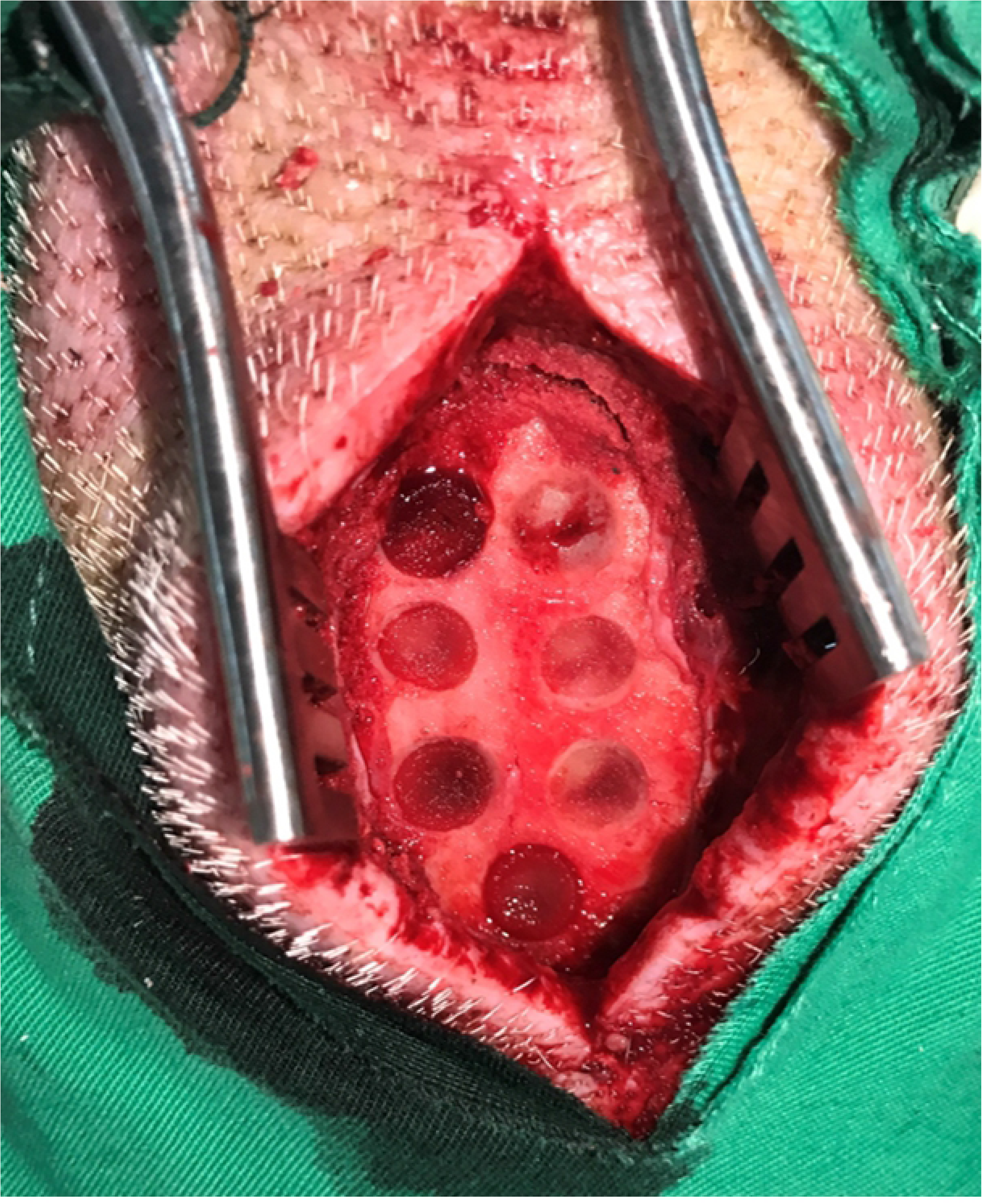
Critical-sized calvarial defects of DsRed pig were created. The frontal bone was exposed after a standard sagittal approach in the forehead region. Seven identical bony defects were created in the frontal bone and the defects were positioned at least 5mm apart to minimize biological interactions. The internal plate of the neurocranium remained completely intact during the procedure. After adequate hemostasis, the scaffolds were implanted.

### Sacrifice and preparation of specimens

Two animals were sacrificed 1, 2, 3, and 4 weeks after transplantation. The animals were anesthetized by intramuscular injection of tiletamine-zolazepam and atropine and then were euthanized with a lethal injection of sodium pentobarbital. The frontal bone was en bloc resected, and the calvarial defects were retrieved separately using an oscillating autopsy saw. The retrieved specimens were fixed in 10% neutral formalin for further survey.

### *In vivo* fluorescence imaging

All samples were washed twice with PBS, fixed in 4% v/v formaldehyde (methanol-free; Polyscience) for 15 min, permeabilized with 0.1% v/v Triton X-100 for 5 min, and incubated in 10 mg/mL bovine serum albumin and 100 g/mL RNase for 45 min at room temperature. F-actin filaments were stained with Alexa Fluor-conjugated phalloidin (Molecular Probes) for 20 min, and nuclei were counterstained with 10 g/mL propidium iodide (Sigma) for 10 min. Finally, the samples were washed with PBS and mounted in Vectashield®. CLSM images were acquired on a BioRad MRC 600 microscope. The quantitative distribution of the two types of fluorescent cells was analyzed by using NIH Image J software (National Institutes of Health, USA) and presented as the modified integrated density (Modified IntDen). The modified integrated density was calculated based on the selected Integrated Density - (Area of selected cell ×Mean fluorescence of background readings).

### Histology

The tissue samples were processed to obtain non-decalcified ground sections. After 2 weeks in 10% neutral formalin, the specimens were rinsed in running tap water, trimmed, dehydrated in ascending concentrations of ethanol, and embedded in methyl methacrylate. The embedded tissue blocks were serially cut into 5-μm-thick ground sections using a microtome (Leica SM 2000; Germany) and mounted on glass slides. Ten cross-sections from the middle of each implant were stained with hematoxylin-eosin and Masson’s trichrome and visualized using an optical microscope (DXM200F Digital Camera; Nikon, Tokyo, Japan).

### Immunohistochemistry of CD68

After fixation in formalin, the dissected blocks were decalcified for 11 days in 10% EDTA and subsequently embedded in paraffin. In brief, after removal of the embedding material, endogenous peroxidases were blocked with H_2_O_2_. Semi-thin sections were rinsed in saline and incubated in phosphate-buffered citrate solution, pH 7.4 (Sigma-Aldrich Corp., St Louis, MO, USA), for 20 min in the microwave and rinsed with phosphate-buffered saline. A microtome was used to obtain 3-mm-thick sections. Immunohistochemistry analysis of tissue labelled with anti-CD68 was performed with purified ab81289 (Abcam Inc., Cambridge, MA) at 1:250. Heat-mediated antigen retrieval was performed using Tris/EDTA buffer pH 9.0 (Abcam Inc., Cambridge, MA). Goat Anti-Rabbit IgG H&L (HRP) was used as the secondary antibody at 1/500. The negative control used PBS instead of the primary antibody, and counterstaining was performed with hematoxylin. The qualitative analysis was performed using a light microscope (Nikon Eclipse E600).

### Statistical analysis

All experimental results are expressed as the mean ± SD, and the statistical analysis was performed using Student’s t-test and two-way ANOVA. Analysis of variance (ANOVA) was performed by using Statistica 6.0 software (Statsoft, Tulsa, OK, USA). Differences with a p value of < 0.05 were considered statistically significant.

## Results

### *In vitro* characterization of EGFP-pMSCs

Passage 5 EGFP-pMSCs [20] were collected by incubation with 0.25% trypsin-EDTA for 5 min at 37°C and re-suspended in ice-cold washing buffer consisting of PBS with 2% FBS. Phase-contrast images of EGFP-MSCs showed a change from a round-shaped to a spindle-shaped morphology after culture for 1 day. After 2 days of culture, 90% confluence was reached, and the cells were trypsinized and re-plated at a dilution of 1:3. These EGFP-pMSCs proliferated rapidly and harbored a greater abundance of EGFP expression by fluorescence microscopy (Fig 3).

**Fig 3.**
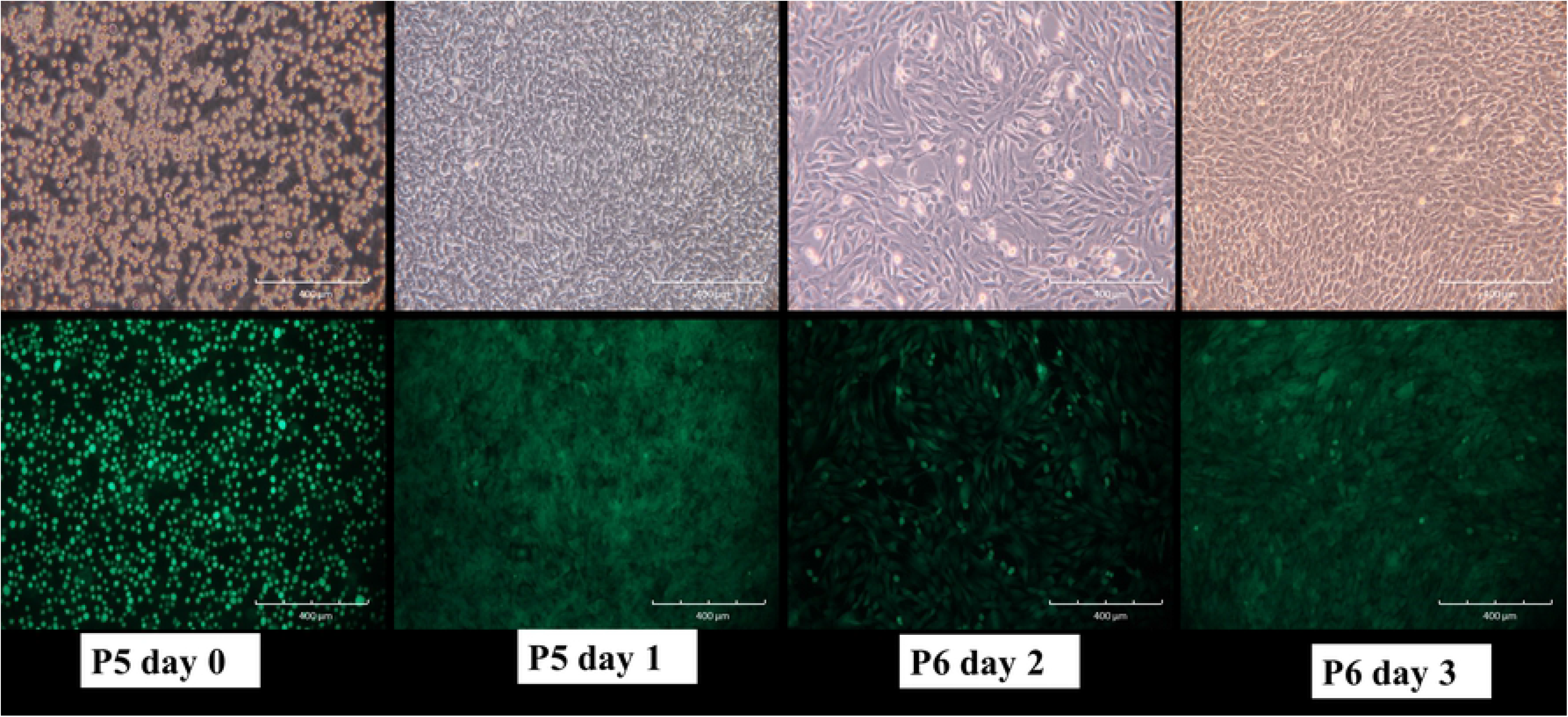
Cell culture of EGFP-MSCs. Phase-contrast image of passage 5 EGFP-MSCs showing the change from round-shaped to spindle-shaped morphology after culture for 1 day (Upper Row). After 2 days of culture, 90% confluence was reached, and the cells were trypsinized and re-plated at a dilution of 1:3. These EGFP-pMSCs proliferated rapidly and harbored a greater abundance of EGFP expression according to fluorescence microscopy (lower row).

### *In vitro* fluorescence evaluation

The sterile scaffolds were individually placed onto the wells of a 48 well-plate. The cell seeding density was 5 × 10^3^ or 5 × 10^4^ cells/scaffold in 50 *μ*L of medium, and the cells were seeded throughout the top side of the scaffold and then incubated for 3 h to allow cell attachment before adding the maintenance medium. From day 3, osteoinduction medium was added, and increasing expression of green fluorescence from day 3 to day 28 was observed in both groups by fluorescent microscopy, especially in the 5 × 10^4^ group (Fig 4).

**Fig 4.**
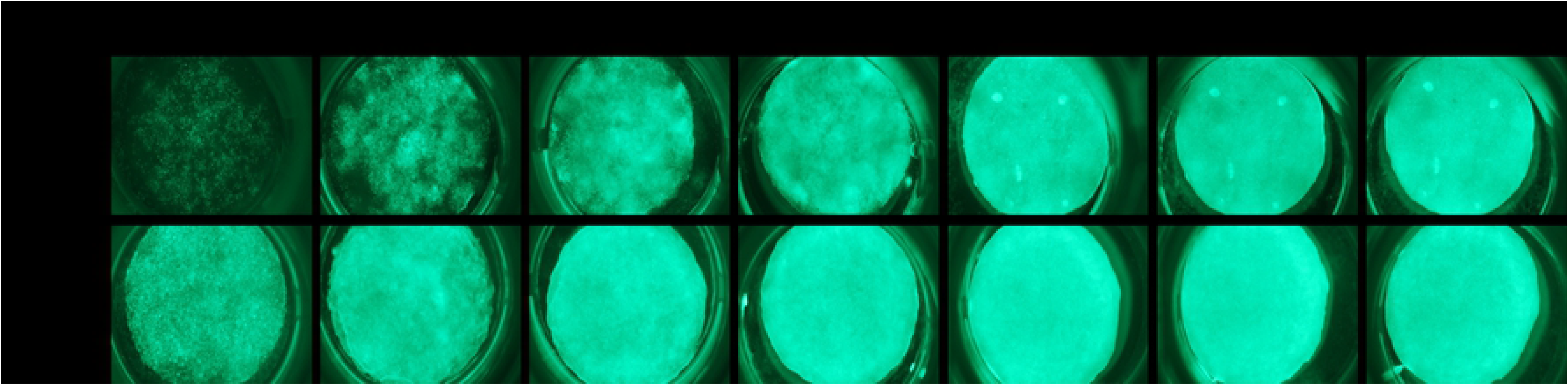
*In vitro* fluorescence evaluation of two different loading density in the scaffolds. An increase in the expression of green fluorescence was observed from day 3 to day 28 in both groups by fluorescence microscopy, especially in the 5 × 10^4^ group.

### *In vitro* cell distribution and morphology analysis

SEM analysis showed that this biodegradable and biocompatible scaffold [36] with a mean pore size of approximately 148 ± 62 μm was suitable for growth of the EGFP-pMSCs. The intensity of live cells increased from day 3 to 28, indicating sufficient intricate space available for ingrowth. SEM images indicated that the EGFP-pMSCs spread on the sponge, and cellular extension and networks were observed. The connected pores were nearly covered by cells by day 21 in the 5×10^4^ group and by day 28 in the 5×10^3^ group (Fig 5). CLSM images of the *in vitro* distribution of EGFP-pMSCs in the scaffolds were obtained from day 3 to day 28 (Fig 6). The constructs were washed with PBS and mounted in Vectashield®. CLSM images were acquired on a BioRad MRC 600 microscope (100X). Three-dimensional z-stack images were evaluated and merged. The distribution of green fluorescence in both groups corresponded to the cell density, especially in the surface view (Fig 6A). In the cross section of the scaffold (Fig 6B), cells were distributed into the scaffold from the surface to the center from day 3 to day 28. On day 28, more cells were observed in the central area in the 5×10^4^ group than the 5×10^3^ group, and the space seemed to be sufficient for cell growth in both groups.

**Fig 5.**
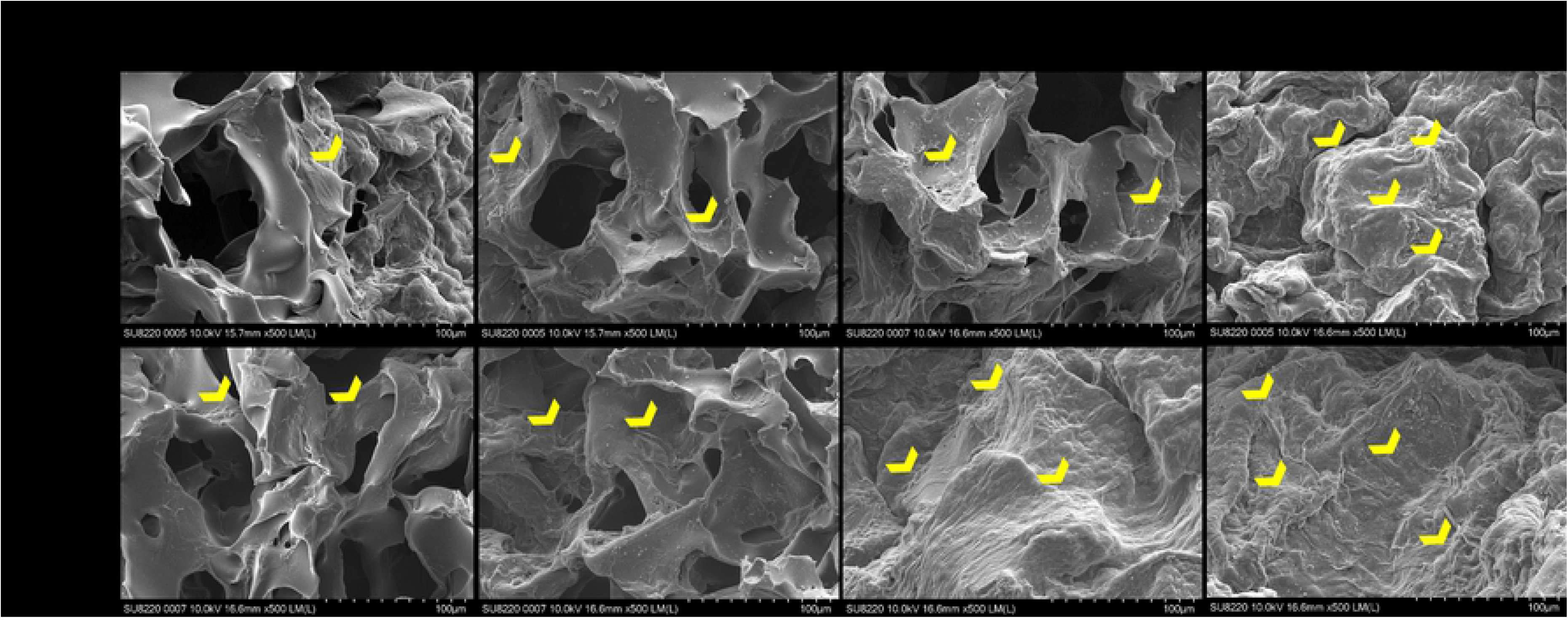
*In vitro* SEM analysis. SEM images indicated that the EGFP-pMSCs (yellow heads) spread on the sponge, and cellular extension and networks between them were observed from day 7 to day 28 in both groups. The connected pore was nearly covered by cells at day 21 in the 5×10^4^ group and day 28 in the 5×10^3^ group.

**Fig 6.**
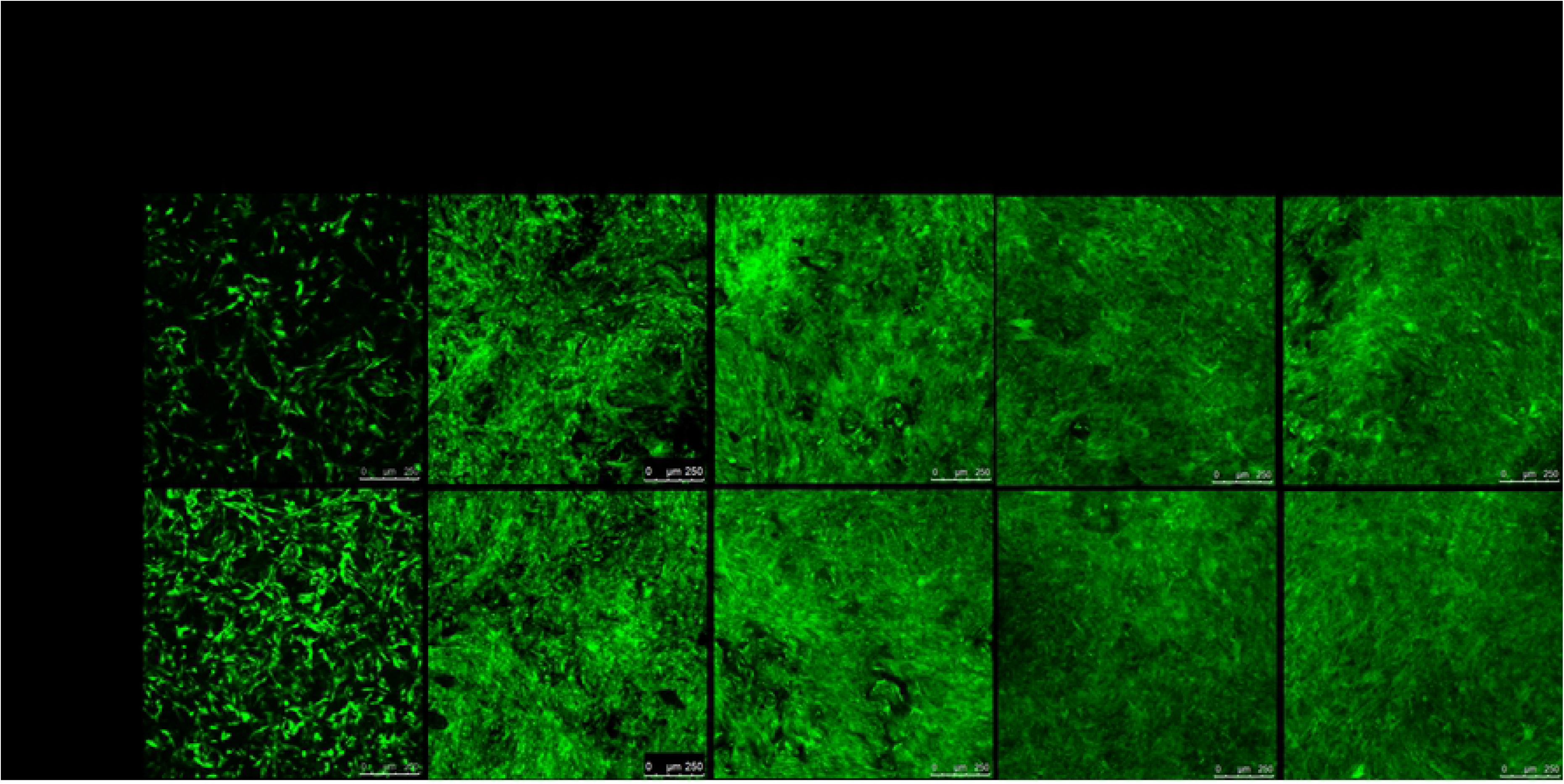
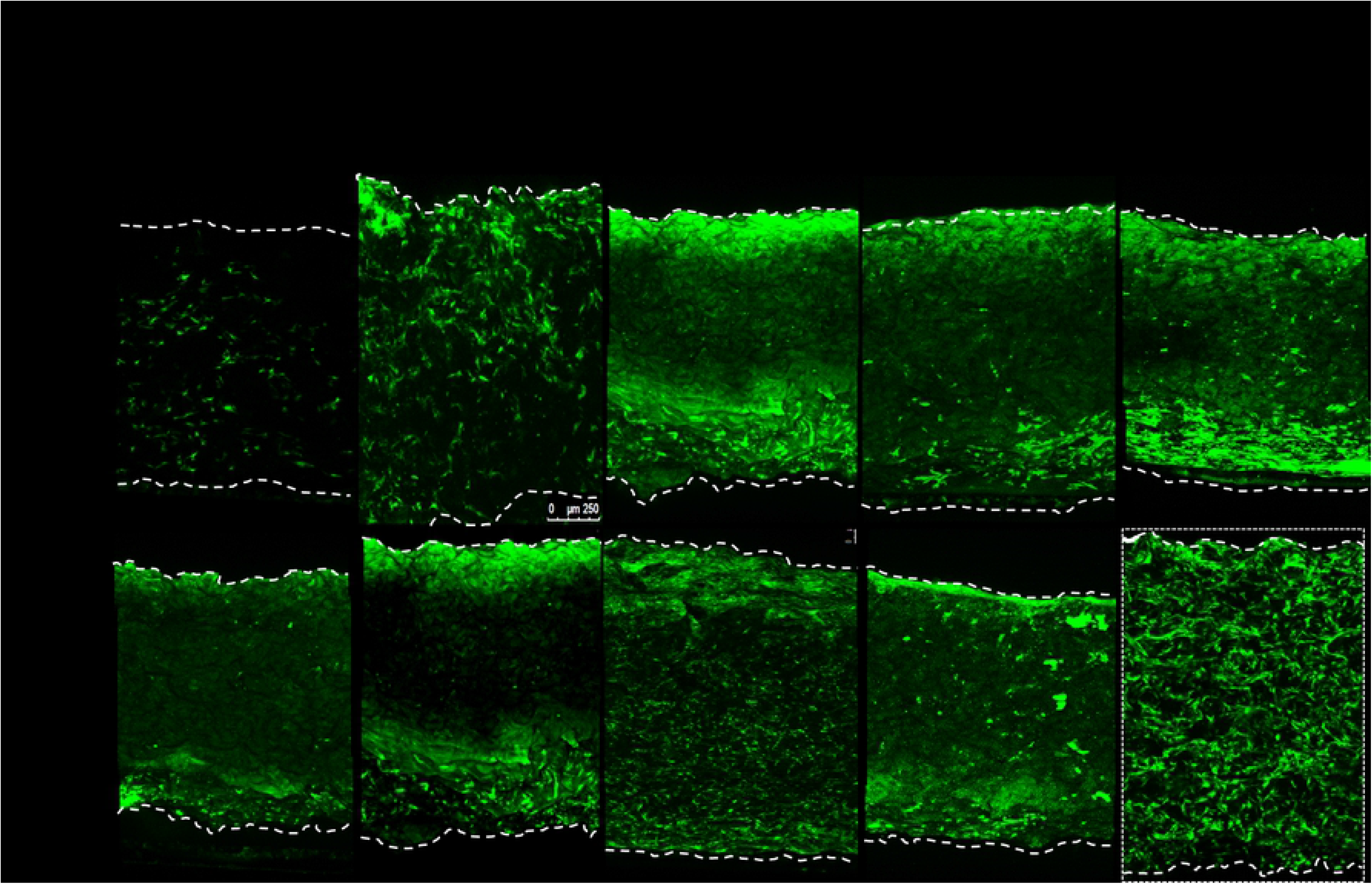
CLSM images of the *in vitro* distribution of EGFP-pMSCs in the scaffolds. The green fluorescence distribution in both groups corresponded to the cell density, especially in the surface view (A). In the cross section of the scaffold (B), cells were distributed into the scaffold from the surface to the center from day 3 to day 28.

### *In vitro* osteogenic differentiation

#### Alkaline phosphatase staining and quantification

The alkaline phosphatase staining results on the culture plates were observed by microscopy (Fig 7A). Both groups showed slight positive staining on day 7, and the relative staining intensity increased on days 14, 21, and 28 compared to days 0 and 7 in the 5×10^3^ cell group. In the 5×10^4^ cell group, the staining intensity increased significantly from day 7. In both groups, the staining intensity continued to slowly increase steadily from day 21 to day 28. Alkaline phosphatase is an early marker of bone formation, and the level increased significantly in the first 2 weeks, followed by a slow steady increase until day 28. Alkaline phosphatase activity (Fig 7B) showed that the cells seeded in the two groups were able to produce alkaline phosphatase on days 7, 14, 21, and 28, and the enzyme activity was significantly higher in the 5×10^4^ group than the 5×10^3^ group between days 7 and 28 (** *p* < 0.01, *** *p* < 0.001), indicating higher osteoblast differentiation in the 5×10^4^ group.

**Fig 7.**
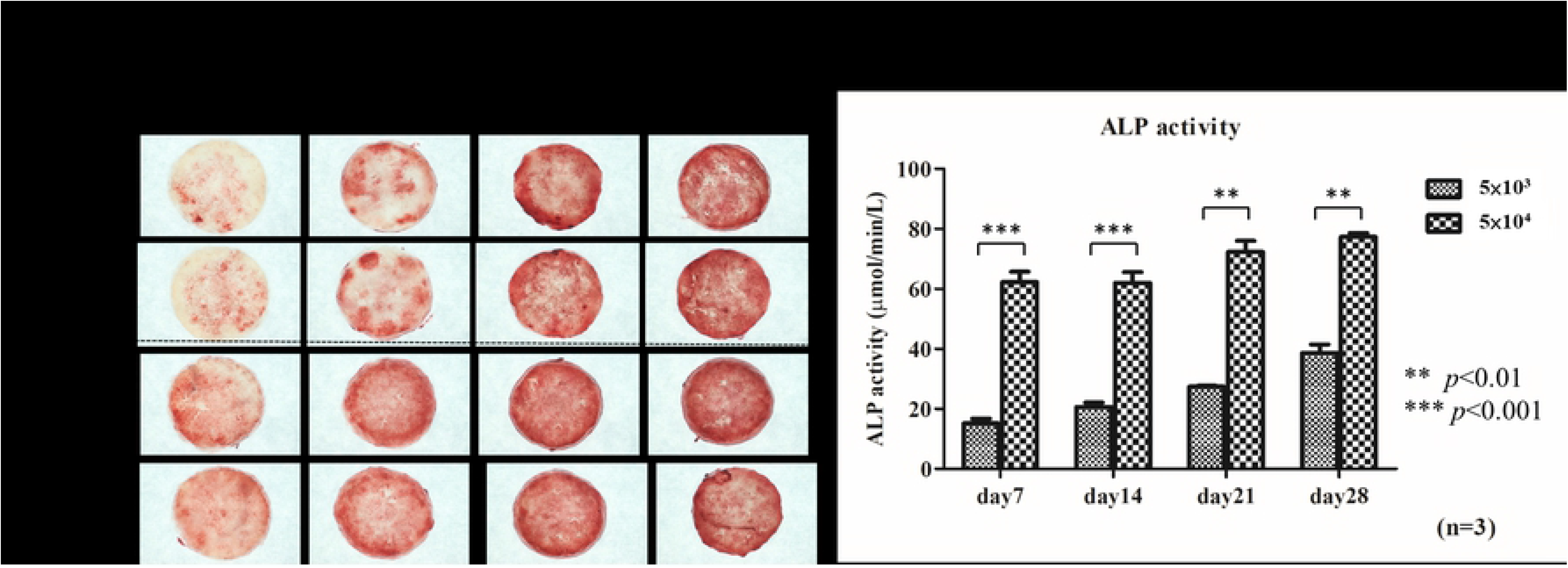
Alkaline phosphatase staining and quantification analysis. Microscopy observation of alkaline phosphatase staining results on the culture plates (A). Both groups showed slight positive staining on day 7, and the relative staining intensity increased on days 14, 21, and 28 compared to Days 0 and 7 in the 5×10^3^ cell group. In the 5×10^4^ cell group, the staining intensity increased significantly from day 7. Alkaline phosphatase activity (B) showed the enzyme activity was significantly higher in the 5×10^4^ group than the 5×10^3^ group from days 7 to 28 (** *p* < 0.01, *** *p* < 0.001), indicating higher osteoblast differentiation in the 5×10^4^ group. Data is shown as mean ± standard deviation.

#### Alizarin red S (ARS) staining and quantification

Alizarin red S staining (Fig 8A) is indicative of bone mineralization and was negative on day 7 in both groups. Up to day 14, all groups showed positive staining with a deep red color, indicating bone nodule formation. The alizarin red staining quantification assay (Fig 8B) was performed at different time points, and the absorbance was measured at 540 nm using an ELISA reader. The absorbance value significantly increased from day 7 to days 14, 21 and 28. The values were significantly higher in the 5×10^4^ group than in the 5×10^3^ group on day 14 and 21 (*p*<0.05), consistent with the results of alkaline phosphatase staining.

**Fig 8.**
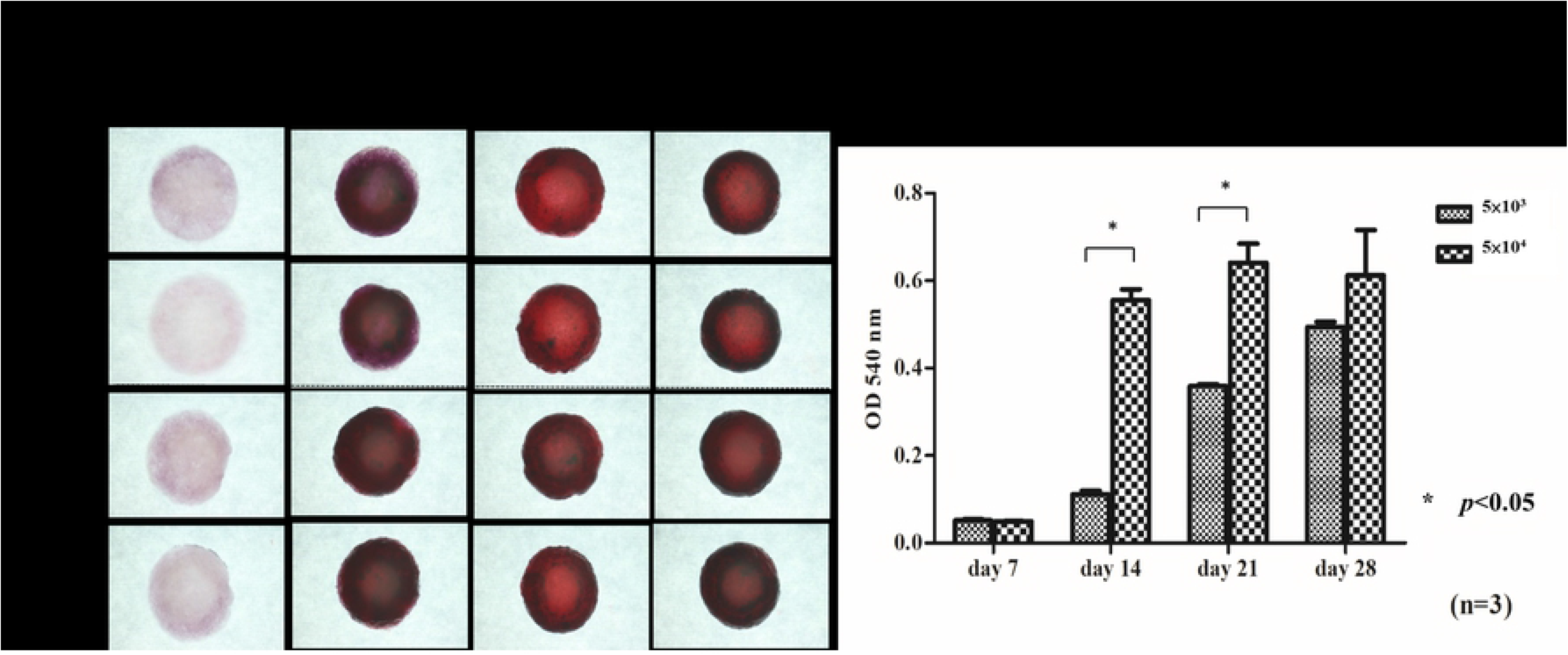
Alizarin red S staining and quantification analysis. Alizarin red S staining (A) was negative on day 7 in both Groups. Up to day 14, all groups showed positive staining with deep red color, indicating bone nodule formation. Alizarin red staining quantification assays (B) were performed at different time points. The absorbance value increased significantly from day 7 to days 14, 21 and 28. The values were significantly higher in the 5×10^4^ group than in the 5×10^3^ group on day 14 and 21. Differences considered significant at * *p*< 0.05 were marked. Data is shown as mean ± standard deviation.

### *In vivo* fluorescence imaging

Confocal laser scanning microscopy was performed to image groups 1 through 5 from week 1 to week 4. (Fig 9). EGFP-pMSCs were present in the scaffolds in groups 4 and 5 through the whole period. Although the EGFP expression decreased in the third and fourth weeks, DsRed expression increased in groups 4 and 5. The green fluorescence was higher in group 5 than in group 4 at each time point, similar to the *in vitro* fluorescence imaging results (Fig 6A). DsRed expression was significantly higher in groups 2 and 3 than in group 1 at each time point, which indicated that the empty scaffold facilitates DsRed cells ingrowth. Even at week 4, no sufficient two fluorescent cells were observed in group 1, which indicates that this type of calvarial defect cannot heal spontaneously during the bone healing period used in our experiments.

**Fig 9.**
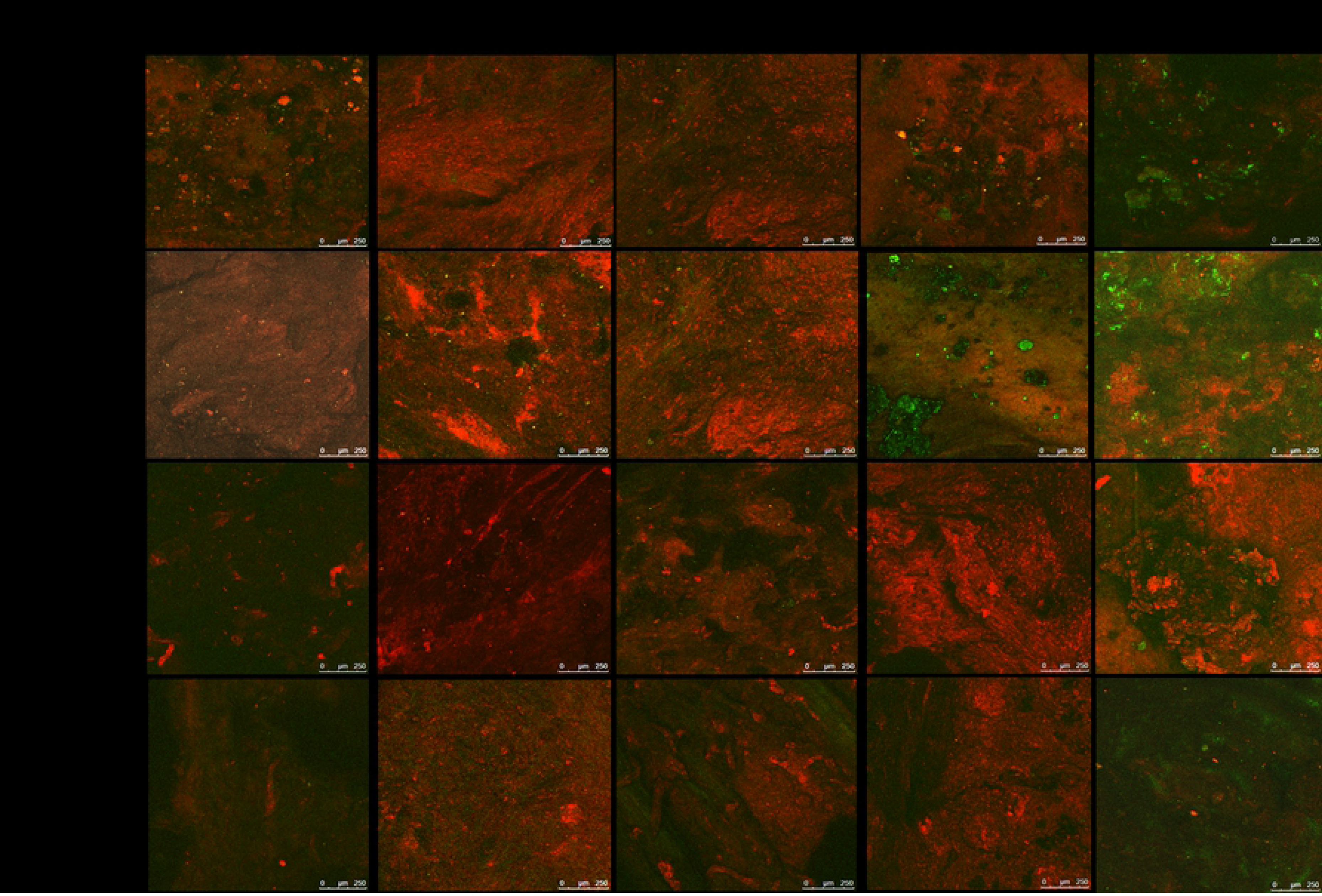
CLSM image of group 1 to 5 from week 1 to week 4. EGFP-pMSCs were present in the scaffolds in groups 4 and 5 from week 1 to week 4. Although EGFP expression decreased in the third and fourth weeks, DsRed expression was increased in groups 4 and 5. Green fluorescence was higher in group 5 than in group 4 at each time point, similar to the *in vitro* fluorescence distribution result. DsRed expression was significantly higher in groups 2 and 3 than in group 1 at each time point, which indicated that the empty scaffold facilitates DsRed cells ingrowth. Even at week 4, no sufficient two fluorescent cells were observed in group 1.

### *In vivo* fluorescence quantification

The modified integrated density of green and DsRed fluorescence in the acquired images was measured using Image J and expressed as bar graphs (Fig 10). EGFP expression (Fig 10A) was higher in Groups 4 and 5 than in the other three groups from the 1^st^ week to the 3^rd^ week (*p*<0.05; *p*<0.01) and was significantly highest in group 5 in the 4^th^ week (*p*<0.05). These results indicate that the seeded cells were present in the scaffold and that the higher load of seeded cells was sustained until the 4^th^ week. The results of the *in vivo* study were compatible with those of the *in vitro* study. By the 4^th^ week, EGFP expression in Group 4 decreased to the levels in Groups 1-3, which means that an insufficient number of seeded cells may not be effective for tissue transplantation. EGFP expression was highest in the 2^nd^ week, which means that the proliferation of seeded cells reached its highest peak, consistent with the *in vitro* ALP/ARS staining results (Figs 7 and 8).

**Fig 10.**
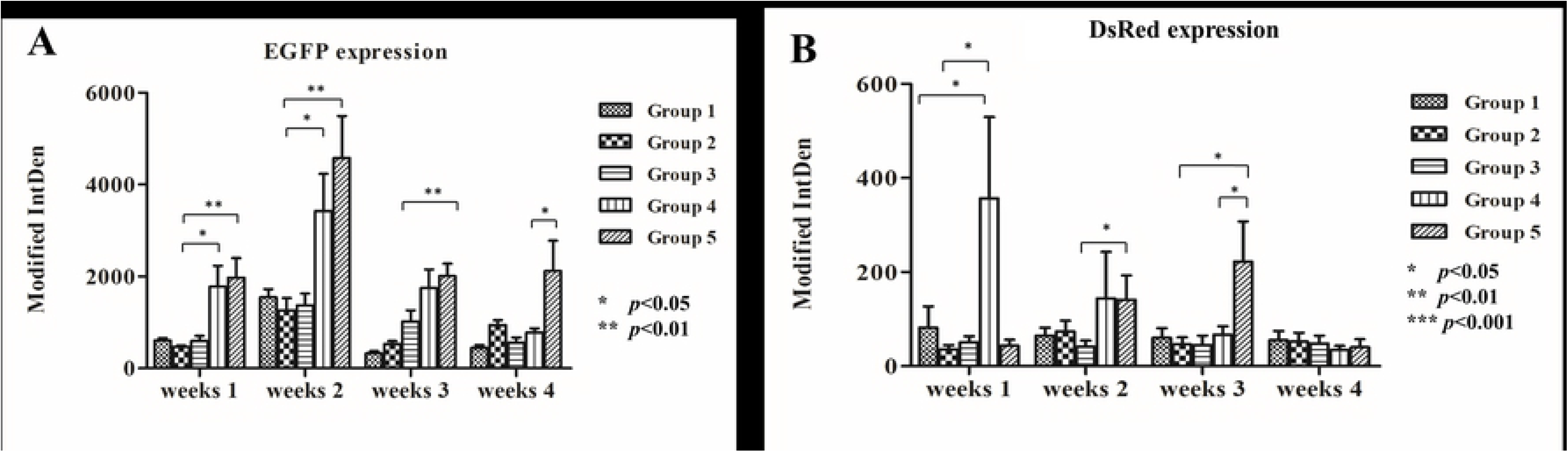
*In vivo* fluorescence quantification. EGFP (A) and DsRed (B) expression using Modified IntDen in Group 1 to 5 from week 1 to 4 were demonstrated as bar chart. Differences considered significant at * *p*< 0.05, ** *p* < 0.01, and *** *p* < 0.001 respect to control. Data is shown as mean ± standard deviation.

In the 1^st^ week, DsRed expression (Fig 10B) was higher in Group 4 than in the other groups, indicating that recruitment of host cells was facilitated by the scaffold seeded with a lower density of cells compared with the empty scaffold or scaffold seeded with a higher density of cells. DsRed expression decreased from the 1^st^ week to 4^th^ week in group 4 but increased in group 5, which means that more EGFP pMSCs can recruit more host cells 2 weeks after transplantation and that enough available space and higher cell loading can benefit tissue regeneration. The DsRed expression in groups 1 to 3 did not increase from the 1^st^ week to 4^th^ week, indicating that the empty scaffold attracted host cells only in the first week after transplantation and implying that scaffolds without seeded cells may not be effective for tissue transplantation. Osteogenic differentiation may be responsible for the decreased expression of EGFP and DsRed expression from week 2 to week 4 in groups 4 and 5.

### Histological analysis

Hematoxylin and eosin staining and Masson’s trichrome staining were analyzed using a light microscope at various levels of magnification.

A greater number of cells were stained with H&E in groups 4 and 5 compared to groups 1-3 in the first week (Figs 11). In the fourth week, obvious deposition of osteoids (yellow arrowhead) was noted in group 5. Calcification of the osteoids led to the formation of primitive trabecular bone in group 5 compared with group 4 in the fourth week. In Groups 1-3, cells were sustained until the fourth week, but no obvious osteoids were found in the 4^th^ week.

**Fig 11.**
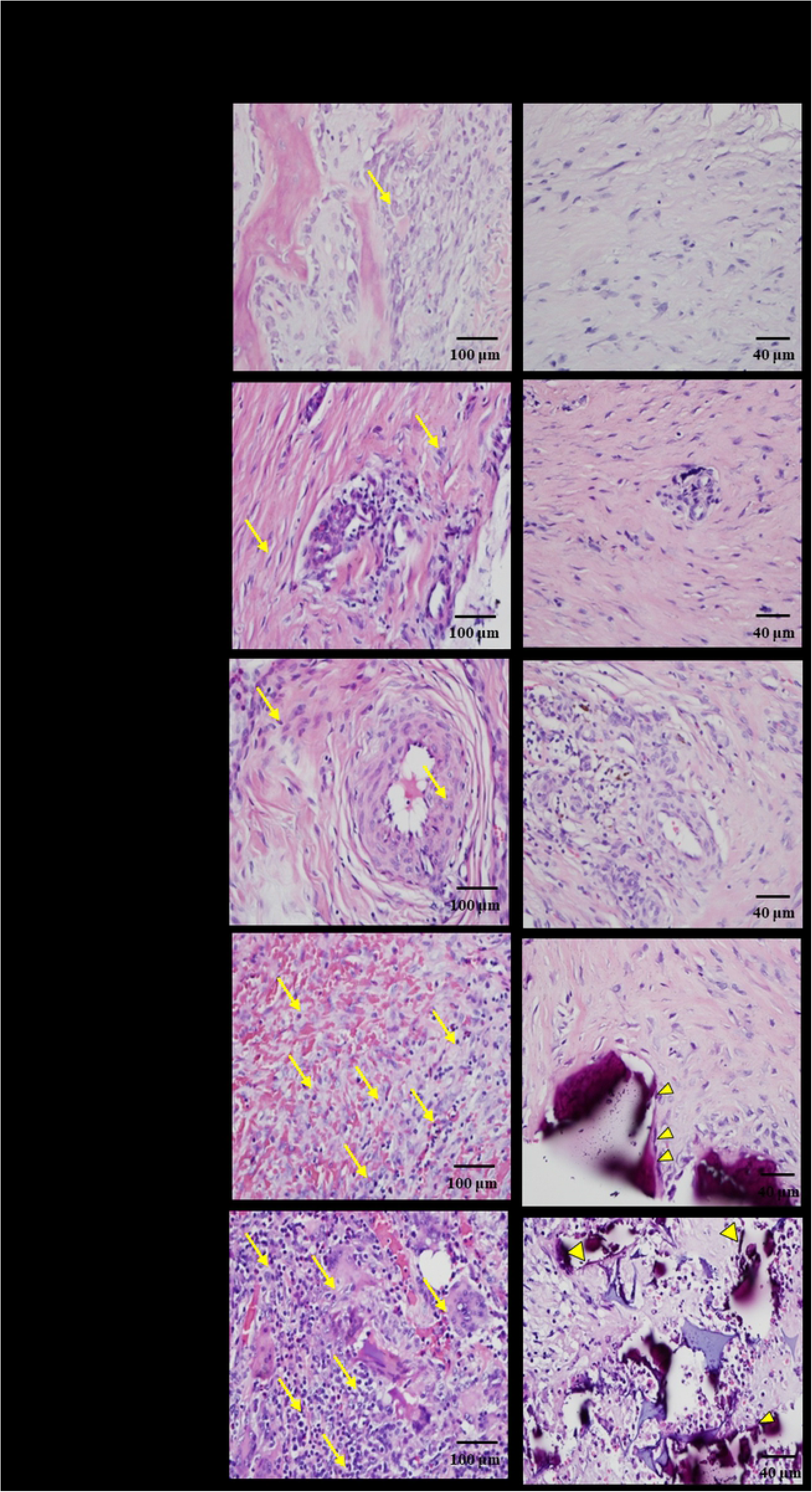
*In vivo* Masson’s trichrome staining. Blue-colored osteoid-like tissue was only found in groups 4 and 5 after the third week. In the 4^th^ week, in groups 4 and 5, thick dense fibrous connective tissue with differentiated osteoblast-like cells (yellow arrow) was noted but only a large amount of undifferentiated cells (yellow arrowhead) without obvious blue-colored osteoid-like tissue in groups 1, 2, and 3. Staining was observed in all groups in 400× magnification.

Masson’s trichrome staining of cells increased in all groups in week 1 through 4, but blue-colored osteoid-like tissue was only found in groups 4 and 5 after the third week (Fig 12). In the 4^th^ week, connective fibrous tissue with a large amount of undifferentiated cells was noted in groups 1, 2, and 3 (yellow arrowhead). Meanwhile, in groups 4 and 5, thick dense fibrous connective tissue with differentiated osteoblast-like cells (yellow arrow) was noted.

**Fig 12.**
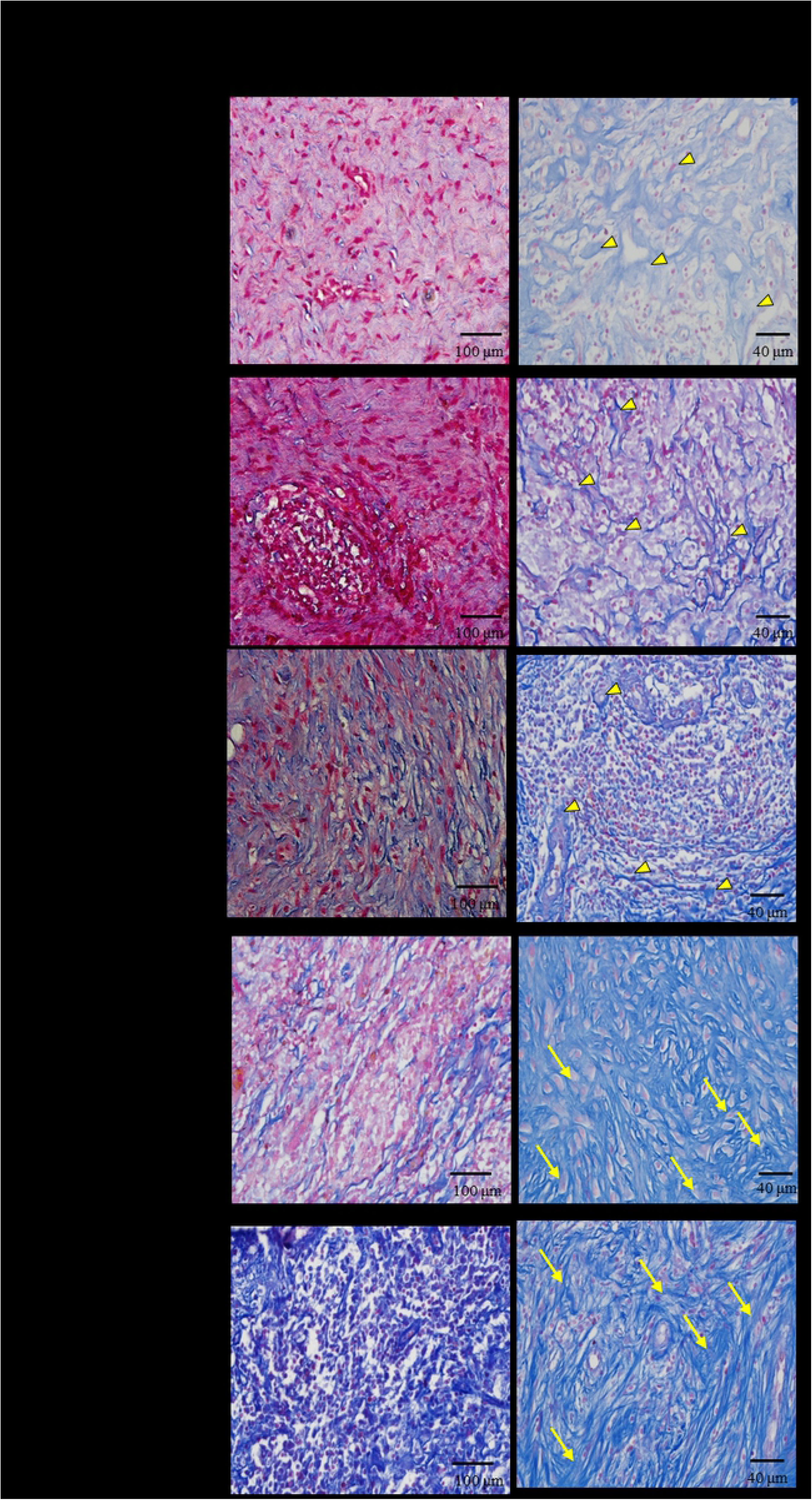
*In vivo* Hematoxylin and eosin staining. A greater number of cells were stained with H&E in groups 4 and 5 compared to groups 1-3 in the first week (yellow arrow). Osteoid formation was confirmed histologically in the hematoxylin and eosin-stained defect of group 4 and 5 at 4 weeks postoperatively (yellow arrowhead). Staining was observed in all groups in 400× magnification.

**Fig 13.**
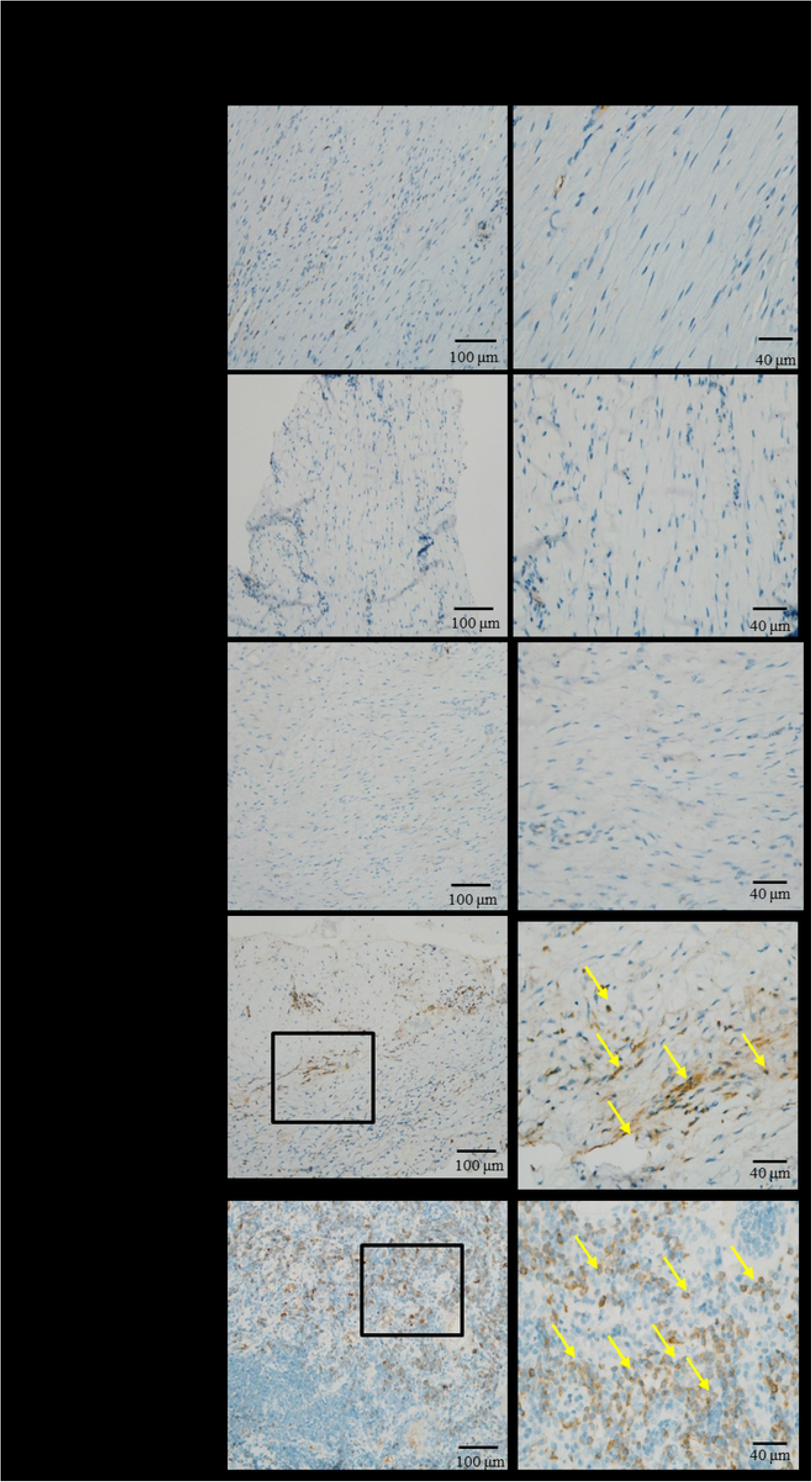
Immunohistochemistry of CD 68. The macrophage marker were stained with anti-CD68 antibody and marked with yellow arrow in group 4 and 5 in week 4. The increasing arrows, CD 68, in group 5 than group 4 suggested that more neovascularization and thus more osteogenesis in group 5 than group 4.

### Immunohistochemistry of CD68

The levels of the macrophage marker [37] CD68 were higher in groups 4 and 5 than in the other groups in the fourth week (Fig 12), especially around the neovascularization area. The increase in macrophages indicated an increase in osteoclasts and, consequently, more osteogenesis in groups 4 and 5.

## Discussion

Allografts may provide the same osteoconductive conduits for bony fusion as traditional autografts and may have comparable biomechanical properties [5, 6]. Although depleted of osteoprogenitor cells like MSCs, the fusion rate still reaches 73% to 100% in instrumented spinal fusion [7-16], making allografts a clinically feasible alternative for fusion. Cells, scaffolds, and factors are the triad of regenerative engineering, and the lack of one of these three graft properties made us wonder about the role of seeded cells in bone regeneration.

A number of studies examining the combination of MSCs and 3-D scaffolds in bone regeneration have reported promising outcomes in animal critical size bone defects [38-43]. These studies aimed to achieve the final result of bone repair by using different kinds of scaffolds [44], sources of MSCs [45] or loaded factors [46-48], but very few studies have focused on the role of the seeded cells.

Kuznetsova et al. [49] used allogenic transplantation of mice MSCs loaded on fabricated 3-D scaffolds made from poly (D, L)-lactic acid and hydroxyapatite in the calvarial defect. They divided the mice into two experimental groups: GFP (+) mice MSCs transplanted into GFP (-) mice and GFP(-) mice MSCs transplanted into GFP(+)mice; the scaffold without cells was used as the control. Fluorescence imaging and histology revealed only allogeneic cells on the scaffold after 6 weeks and 12 weeks, and newly formed bone-like tissue from seeded cells was detected by 12 weeks. The presence of vessels was confirmed by CD31 immunohistochemical staining, which indicated the possibility of vessel formation from seeded MSCs.

However, there were four major drawbacks of their study. First, the use of Hoechst stain in fluorescence imaging to indicate seeded or host cells is not reasonable because the GFP harvest of cells may decay after transplantation for 6 weeks or 12 weeks; cells with GFP may also stain with blue color, providing misleading results about the origin of cells. Second, the surface view of fluorescence imaging cannot represent the actual cell interactions during bone regeneration inside the 3-D scaffold. Third, the creation of only one defect in each mouse cannot eliminate individual variations. Fourth, bone formation was supposed to be similar in both experimental groups. The authors explained that GFP itself may be immunogenic in immunocompetent mice; however, GFP was present in both groups.

To the best of the authors’ knowledge, the present study is the first to compare the distribution and proportion of seeded cells and host cells by tracking two fluorescent cells in the same scaffold in a pig critical-sized calvarial defect model.

To eliminate individual variations in the animal study, we used a critical-sized calvarial defect in pigs instead of rats due to the limited size of rats. As an animal model, we chose the domestic pig because its bone regeneration rates correlate well with those found in humans (pigs 1.2 −1.5 mm per day, humans 1.0 - 1.5 mm per day) [34]. Numerous experiments have been conducted to analyze bone formation with various scaffolds and factors in critical-sized calvarial defect pigs [50-52] because of the uniform, flat and surgically feasible calvarial bone. In general, a defect size with a diameter of at least 6mm is thought to be a reasonable critical-sized calvarial defect, and we also used an empty group (group 1) as a control.

The CAG hybrid promoter-driven EGFP-MSCs derived from the transgenic pigs used in this study have been proved to exhibit homogeneous surface epitopes and possess classic trilineage differentiation potential into osteogenic, adipogenic, and chondrogenic lineages, with robust EGFP expression maintained in all differentiated progeny. These cells have the major advantage of greater sustained EGFP expression in stem cells and in all their differentiated progeny [19] compared to GFP expression in transgenic pigs under the control of housekeeping regulatory sequences [53,54]. Increased expression of green fluorescence was observed from day 3 to day 28 after culture into scaffolds (Fig 4.and 6)

The transgenic Ds-Red pigs were produced by using pronuclear microinjection, and the transgene comprised a CMV enhancer/chicken-beta actin promoter and DsRed monomeric cDNA [33]. DsRed is a red fluorescent protein that was originally discovered in corals of the class Anthozoa. DsRed is expressed ubiquitously, with red fluorescence detected in the brain, eye, tongue, heart, lung, liver, pancreas, spleen, stomach, small intestine, large intestine, kidney, testis, and muscle, as confirmed by histology and western blot analyses. Red fluorescence tracking was previously applied in transplantation of neurons from Ds-Red transgenic pigs into a Parkinson’s disease rat model [55]. In this *in vivo* experiment, precise evaluation and analysis could be achieved due to the abundant expression of the two fluorescent proteins in all differentiated progeny.

The hemostatic gelatin sponge, Spongostan^TM^, is prepared from purified type A pork skin, and its biosafety in clinical applications, especially to control bleeding during surgery, has long been proved [56, 57]. *In vivo* studies have shown good biocompatibility of Spongostan^TM^ without any immune response during bone formation [36]. The main reasons that we chose this scaffold are its biocompatibility and biodegradation with MSCs and absence of autofluorescence [36].

Our *in vivo* fluorescence analysis (Fig 9 and 10) showed that the green fluorescence in the scaffold loaded with seeded cells in groups 4 and 5 was significantly higher than that in the other groups from week 1 to week 4, which means that the seeded cells did play an important role in the osteogenic differentiation process. EGFP-pMSCs were highest in group 5 in the 4^th^ week, which means that the higher load of seeded cells was sustained until the 4^th^ week. DsRed expression (Fig10) was highest in Group 4 in the 1^st^ week, indicating that recruitment [58, 59] of host cells was facilitated in the lower density-seeded scaffold than the empty scaffold or higher density-seeded scaffold. DsRed expression decreased from the 1^st^ week to 4^th^ week in group 4 but increased in group 5, which means that more EGFP-pMSCs can recruit more host cells in the following period and that enough available space and higher cell loading benefit tissue regeneration. The higher expression of EGFP and DsRed fluorescence in groups 4 and 5 were compatible with the histological results, which indicated more obvious deposition of osteoids in groups 4 and 5 in week 4. The immunohistochemical results for CD68 in week 4 also showed obvious osteogenesis in groups 4 and 5.

Our *in vivo* results demonstrated that the seeded MSCs promoted host cell migration, which contributed to the higher number of host cells in the transplanted scaffolds. In addition, according to the histological staining, we found that the percentage of osteoid formation in the scaffolds transplanted with seeded MSCs was significantly higher than that in the control group, which means that not only the transplanted seeded cells but also the recruited host cells contributed to the entire process.

In summary, we found that more seeded cells recruit more host cells and that both cell types participate in osteogenesis. Recruitment of host cells was facilitated in the lower density-seeded scaffold than the empty scaffold or higher density-seeded scaffold in the first week but sustained recruitment of host cells was achieved in higher density-seeded scaffold and result in more obvious deposition of osteoids. Scaffolds without seeded cells may therefore not be effective in bone transplantation based on this *in vivo* study.

## Acknowledgments

This work was supported by the Chang Gung Memorial Hospital (grant No. CMRPG3G2061; CMRPG3G0572). We would like to thank the Microscope Core Laboratory at the Center for Advanced Molecular Imaging and Translation and the Laboratory Animal Center of Linko Chang Gung Memorial Hospital.

